# Adjuvant selection for optimally balanced humoral and cellular immunity induced by SARS-CoV-2 Spike virosome vaccines

**DOI:** 10.64898/2026.06.23.733553

**Authors:** Marloes Grobben, Gius Kerster, Esther Siteur-van Rijnstra, Mitch Brinkkemper, Meliawati Poniman, Judith A. Burger, Khadija Tejjani, Jacqueline van Rijswijk, Widad Ait Addouch, Melissa Oomen, Joey H. Bouhuijs, Tom Bijl, Ronald Kempers, Kwinten Sliepen, Toon Stegmann, Marit J. van Gils, Mathieu Claireaux, Yme U. van der Velden, Rogier W. Sanders

## Abstract

Current SARS-CoV-2 vaccines provide limited breadth of protection, underscoring the need for vaccine strategies that optimize immune responses. Virosomes offer a modular vaccine platform that enables multivalent antigen display and incorporation of adjuvants which can steer immune responses. We evaluated the immune response in BALB/c mice with virosomes displaying SARS-CoV-2 Wuhan or Delta spike antigens and coupled with various distinct adjuvants. Adjuvant selection differentially influenced both humoral and cellular immune outcomes. The TLR7/8 agonist 3M −052 induced a strong Th1-biased response, characterized by elevated IgG2a/IgG1 ratios and robust type 1 cytokine induction with suppression of Th2-associated cytokines. In contrast, the saponin QS-21 enhanced antibody functional quality, illustrated by improved virus neutralization potency and breadth. Furthermore, the combined incorporation of both 3M-052 and QS-21 induced an elevated Th1-biased response without improving neutralization capacity. In conclusion, different adjuvants added onto our virosome-based vaccine led to distinct antibody responses and splenic T-cell profiles, reflective of differences in immune programming. This information guides the selection of adjuvants for respiratory virus vaccines.

## Introduction

SARS-CoV-2, the causative agent of the COVID-19 pandemic, has infected close to a billion people since its emergence (World Health Organization, 2023). The fast-track development and rollout of COVID-19 vaccination programs has saved millions of lives, but efficacy of current vaccines is relatively short-lasting and requires booster shots to keep up immunity (Doria-Rose et al., 2021; Pegu et al., 2021; Suthar et al., 2022). Moreover, the continued emergence of antigenically distinct viral variants compromises cross-protective immunity elicited by first-generation vaccines (Jeworowski et al., 2024). These challenges highlight the need for next-generation vaccine strategies that not only increase antibody breadth but also promote functional humoral and cellular responses.

Virosomes represent a versatile and clinically validated vaccine platform that offers several advantages over soluble protein-based vaccines, including efficient lymphoid trafficking, modular antigen display, and enhanced avidity through multimeric antigen presentation. Virosomes are non-replicating lipid nanoparticles, approximately 100-150 nm in size, composed of viral envelope glycoproteins and synthetic lipids, and lack viral nucleic acids. Influenza-derived virosomes are reconstituted from hemagglutinin (HA, subtype 1), neuraminidase (NA, subtype 1), and synthetic lipids (Moser et al., 2007). Advances in click-chemistry-based conjugation enable rapid and stable attachment of heterologous antigens, such as the SARS-CoV-2 spike protein, to virosomes. Multivalent antigen display on virosomes enhances B cell receptor engagement and promotes efficient antigen delivery to draining lymph nodes, thereby supporting robust humoral immune responses (Chattopadhyay et al., 2017; Wamhoff et al., 2024). The HA and NA components of influenza virosomes offer additional T cell help through intrastructural help whereby pre-existing immunity against influenza glycoproteins may aid de-novo anti-Spike immune responses (Damm et al., 2023; Elsayed et al., 2018)

Currently, there are two successful licensed virosome-based vaccines on the market, Epaxal® (hepatitis A virus) and Inflexal® (Influenza virus; (Moser et al., 2011). Virosome vaccines against HIV-1 (MYM-V201) and malaria (PEV301 & PEV302) are in clinical phase testing (Genton et al., 2007; Lakhashe et al., 2022). More recently, influenza virosomes coupled with SARS-CoV-2 RBD and adjuvanted with 3M-052 successfully protected macaques against severe COVID-19 (Koopman et al., 2023). Additionally, we have previously shown an effective induction of neutralizing antibodies against SARS-CoV-2 using this virosome vaccine platform in mice (Van Der Velden et al., 2022).

Beyond antigen display, adjuvant selection is a critical determinant of immune quality and polarization, particularly for respiratory viral infections. The balance between Th1- and Th2-associated immune responses has important implications for both protection and safety. Th2-skewed immune responses have been associated with immunopathology and vaccine-associated enhanced respiratory disease (VAERD), whereas Th1-biased responses correlate with improved viral control and protective immunity (Zhao et al., 2023). Clinical studies have reported higher mortality in COVID-19 patients exhibiting strongly Th2-polarized immune profiles (Pavel et al., 2021), and preclinical vaccine studies on SARS-CoV-1 have demonstrated disease enhancement following Th2-skewed responses (Tseng et al., 2012). Furthermore, a RSV vaccine formulated with Th1-polarizing adjuvants such as MPLA prevented disease enhancement in murine models (Kamphuis et al., 2013). Therefore, modulation of the type of immune response elicited by vaccines can be beneficial depending on the target pathogen.

Virosomes act as a self-adjuvant by modulating cellular responses through HA mediated cell fusion and subsequent digestion and presentation of antigenic peptides by host cells (Huckriede et al., 2005). The addition of adjuvants to the virosome gives us the ability to tailor the immune responses towards a favorable profile. Several clinically relevant adjuvants have been shown to modulate immune polarization through distinct innate immune pathways. The TLR7/8 agonist 3M-052 induces potent type I interferon signaling and Th1-associated immune responses, the TLR4 agonists MPLA and 3D-PHAD promote innate activation with more moderate polarization towards Th1, and the inflammasome-activating saponin QS-21 induces a balanced Th1 and Th2 profile and enhances T cell activation and antibody functionality (Hernandez et al., 2019; Talukdar et al., 2025). QS-21 has been shown to enhance antibody affinity maturation and functional quality in multiple vaccine platforms (Didierlaurent et al., 2014; Garçon et al., 2012; Talukdar et al., 2025). However, while these adjuvants have been extensively studied in soluble protein- or alum-adjuvanted vaccines, their comparative effects when incorporated into virosome-based vaccine platforms remain poorly defined.

Here, we evaluate influenza-derived virosomes displaying SARS-CoV-2 Wuhan or Delta spike antigens in combination with relevant adjuvants in pseudo influenza-experienced mice. By integrating analyses of antibody magnitude, IgG subclass distribution, neutralization breadth, antigen-specific cytokine secretion, and multivariate modeling, we demonstrate that adjuvantselection differentially shapes immune quality and polarization. Our findings reveal distinct immunological pathways driven by individual adjuvants, with 3M-052 promoting strong Th1 polarization and IgG2a subclass skewing and QS-21 promoting a balanced Th1 and Th2 humoral and cellular immune signature and enhanced antibody functional quality including cross-neutralization. A combination of adjuvants e.g. 3M-052 plus QS-21, however, showed no synergistic effects between IgG2a skewing and enhanced neutralization capacity. These results underscore the importance of rational adjuvant selection for the virosome-based vaccine platform and provide insights into vaccines targeting respiratory viruses.

## Results

### Adjuvant formulation determines the magnitude and Th1/Th2 bias of SARS-CoV-2 spike-specific antibody responses

BALB/c mice were immunized with influenza-derived virosomes displaying the Wuhan spike protein and formulated with distinct adjuvants, to determine how adjuvant formulation influences the magnitude and Th1/Th2 bias of antibody responses elicited by SARS-CoV-2 virosome vaccines (Fig. 1A). Virosomes were prepared with either the TLR7/8 agonist 3M-052, the TLR4 agonist 3D-PHAD, or the saponin QS-21, as mono-adjuvant formulations, or with a combination of QS-21 and 3D-PHAD, alongside non-adjuvanted virosomes as a comparison group. Mice were primed with uncoupled bare influenza-derived virosomes at week 0, to model pre-existing influenza immunity present in humans. Animals (n = 12 per group) received a prime-boost regimen (weeks 3 and 6), and antigen-specific antibody responses were assessed longitudinally in serum and in bronchoalveolar lavage fluid at week 8 (Fig. 1B). Mice were euthanized at week 13. Induction of plasma anti-Wuhan spike IgG was quantified using a Luminex binding assay. All immunized mice developed robust spike-specific IgG responses, which declined between week 8 and 13 (Fig. S1). As expected, prime and booster immunizations also increased anti-hemagglutinin (HA) antibody levels across all groups, reflecting the immunogenicity of the HA component of the virosomes (Fig. S1A).

**Figure 1.**
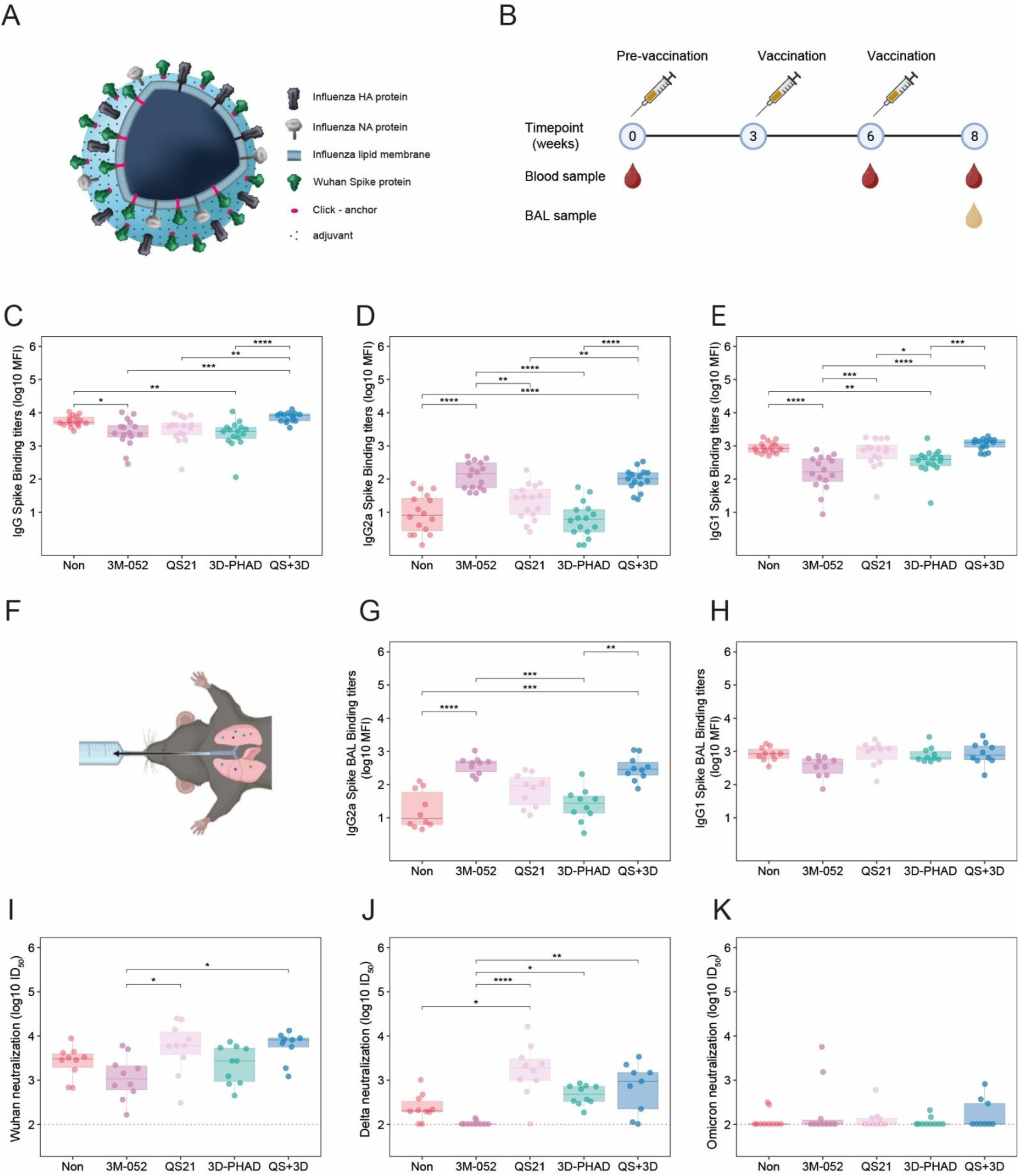
Adjuvant dependent antibody and neutralizing responses. BALB/c mice were immunized with SARS-CoV-2 Wuhan Spike virosomes conjugated with the following adjuvants: non-adjuvanted, 3M-052, QS21, 3D-PHAD or QS21+3D-PHAD. (**A**) An illustration of adjuvant-conjugated Wuhan Spike virosomes. (**B**) Serum and BAL samples were collected at indicated timepoints. ( **C-H**) Antigen-specific total IgG, IgG1 and IgG2a responses at week 8 were quantified for serum (**C-E**) and for bronchoalveolar lavage (BAL) (**F-H**) using a Luminex assay and reported as log10 median fluorescence intensity (MFI). (**I-K**) Neutralizing activity was measured using a pseudotyped virus neutralization assay reported as log10 ID50 against Wuhan (**I**), Delta (**J**) and Omicron (**K**). Data are shown for n = 16 mice per adjuvant group. Statistical comparisons were performed using the Kruskal-Wallis test with Benjamini-Hochberg (BH) correction for multiple comparisons. Dotted lines represent the lower limit of detection. * = P < 0.05, ** = P < 0.01, *** = P < 0.001, **** = P <0.0001.

At week 8, spike-specific IgG levels in mice vaccinated with QS-21- or QS-21 + 3D-PHAD-adjuvanted virosomes were comparable to those observed in mice receiving non-adjuvanted virosomes (median log10 MFI 3.58 and 3.70, respectively) (Fig. 1C). In contrast, mice vaccinated with 3M-052- or 3D-PHAD-adjuvanted virosomes exhibited significantly lower plasma spike-specific IgG levels compared to the non-adjuvanted group (median log10 MFI 3.37 and 3.42, respectively; *P* < 0.05). To evaluate adjuvant-driven antibody isotype polarization, we further quantified this IgG response into anti-spike IgG2a and IgG1 plasma responses following immunization. Mice adjuvanted with 3M-052 induced the highest levels of anti-Wuhan spike IgG2a (median log10 MFI 2.15), exceeding those observed in unadjuvanted virosomes (0.90) and single-adjuvant formulations (QS21, 1.44; 3D-PHAD, 0.78) (Fig. 1D). Notably, the combined QS21 + 3D-PHAD formulation elicited IgG2a levels comparable to 3M-052 (median MFI 2.00). In contrast, 3M-052-adjuvanted mice exhibited lower IgG1 responses (median log10 MFI 2.21), with a similar trend observed for 3D-PHAD (2.58), relative to non-adjuvanted controls (2.92) (Fig. 1E).

Given the reduced total anti-spike IgG levels observed in some adjuvant groups, IgG1 and IgG2a MFIs were normalized to total IgG to assess relative isotype contributions. Following normalization, IgG1 reduction was observed only in the 3M-052 adjuvant group, whereas the IgG1 reduction in the 3D-PHAD group was no longer evident, indicating that the reduced IgG1 level in the 3D-PHAD group was primarily caused by a lower overall anti-Wuhan IgG antibody secretion rather than an IgG subset redistribution (Fig. S1B). This is further confirmed by the lack of difference in ratio of IgG2a over IgG1 levels between 3D-PHAD and non-adjuvanted group of mice. 3M-052 and QS21+3D-PHAD adjuvanted mice elicited a stronger Th1 polarization, reflected by higher IgG2a/IgG1 ratios, compared to non-adjuvanted mice (Fig. S1D). In contrast, unadjuvanted, QS21- and 3D-PHAD-adjuvanted mice exhibited a predominantly Th2-skewed response, reflected by low IgG2a/IgG1 ratios in serum.

As the lung represents the primary site of SARS-CoV-2 infection, bronchoalveolar lavage (BAL) fluid was collected at the time of sacrifice and anti-Wuhan spike-specific antibodies were measured. BAL antibody responses mirrored those observed in serum, with no differences detected in IgG1 levels across groups, whereas 3M-052 and QS+3D induced higher IgG2a titers (Fig. 1G-H, Fig. S1C). A Th2-skewed response was also observed in BAL in unadjuvanted, QS21- and 3D-PHAD-adjuvanted mice while 3M-052 and QS21+3D-PHAD adjuvanted groups showed a more Th1 dominant antibody response also in BAL (Fig. S1E).

### QS-21 enhances the quality and breadth of neutralizing antibody responses

We next assessed the neutralizing capacity of plasma antibodies after vaccination using a pseudovirus-based neutralization assay. All groups showed autologous neutralization against the Wuhan strain, however none of the adjuvanted formulations showed a significant increase in autologous neutralization titers compared to non - adjuvanted virosomes (Fig. 1I).

To evaluate neutralization breadth, heterologous neutralization against the Delta and Omicron BA.1 variants was assessed. Delta cross-neutralization was significantly higher in QS-21-adjuvanted mice compared to non-adjuvanted controls (median log10 ID50 3.27 vs. 2.30, P < 0.05). In contrast, 3M-052-adjuvanted mice exhibited significantly lower Delta neutralization compared to all other adjuvant groups (median log10 ID50 2.00 vs. 3.27, 2.67, and 2.97 for QS-21, 3D-PHAD, and QS-21 + 3D-PHAD, respectively; P < 0.0001, P < 0.05, and P < 0.01) (Fig. 1J). Neutralization of the Omicron BA.1 variant was mostly undetectable across all groups (Fig. 1K).

Because neutralization potency is influenced by antibody abundance, we next assessed antibody quality by normalizing neutralization titers to levels of anti-Wuhan spike IgG (Fig. S1F). After normalization, QS-21-adjuvanted virosomes induced antibodies with significantly improved autologous neutralizing activity (P < 0.01; Fig. S 1F) and enhanced Delta cross-neutralization (P < 0.01; Fig. S1G) compared to the non-adjuvanted formulation. Collectively, these data indicate that QS-21 most effectively enhances neutralization potencyand breadth, whereasother adjuvants provide limited or no improvement.

### 3M-052 induces Th1-biased cytokine secretion while QS-21 promotes polyfunctional T cells

Adjuvantation can differentially shape CD4+ T cell responses which in turn influences the magnitude and quality of antibody responses. Therefore, we assessed antigen-specific T-cell responses by stimulating splenocytes from immunized mice with Wuhan spike antigens and quantified cytokine secretion using a Luminex assay (Fig. 2A). Distinct cytokine profiles were observed across the adjuvant groups which can be stratified into two groups namely Th1 associated cytokines such as IFN-y, TNF-α and IL-2, and Th2 associated cytokines such as IL-4, IL-5 and IL-13. GM-CSF and IL-10 levels were also measured. Splenocytes from mice immunized with 3M-052-adjuvanted virosomes produced high levels of Th1-associated cytokines, including IFN-γ and TNF-α, whereas type 2 cytokines (IL-4, IL-5, and IL-13) were reduced to near undetectable levels compared with the non-adjuvanted group (P < 0.05, P < 0.001, and P < 0.001, respectively). IL-10 secretion was likewise markedly reduced in the 3M-052 group. In contrast, QS-21-adjuvanted virosomes induced the highest levels of GM-CSF and IL-2 among all groups albeit not statistically significantly different from the non-adjuvanted group.

**Figure 2.**
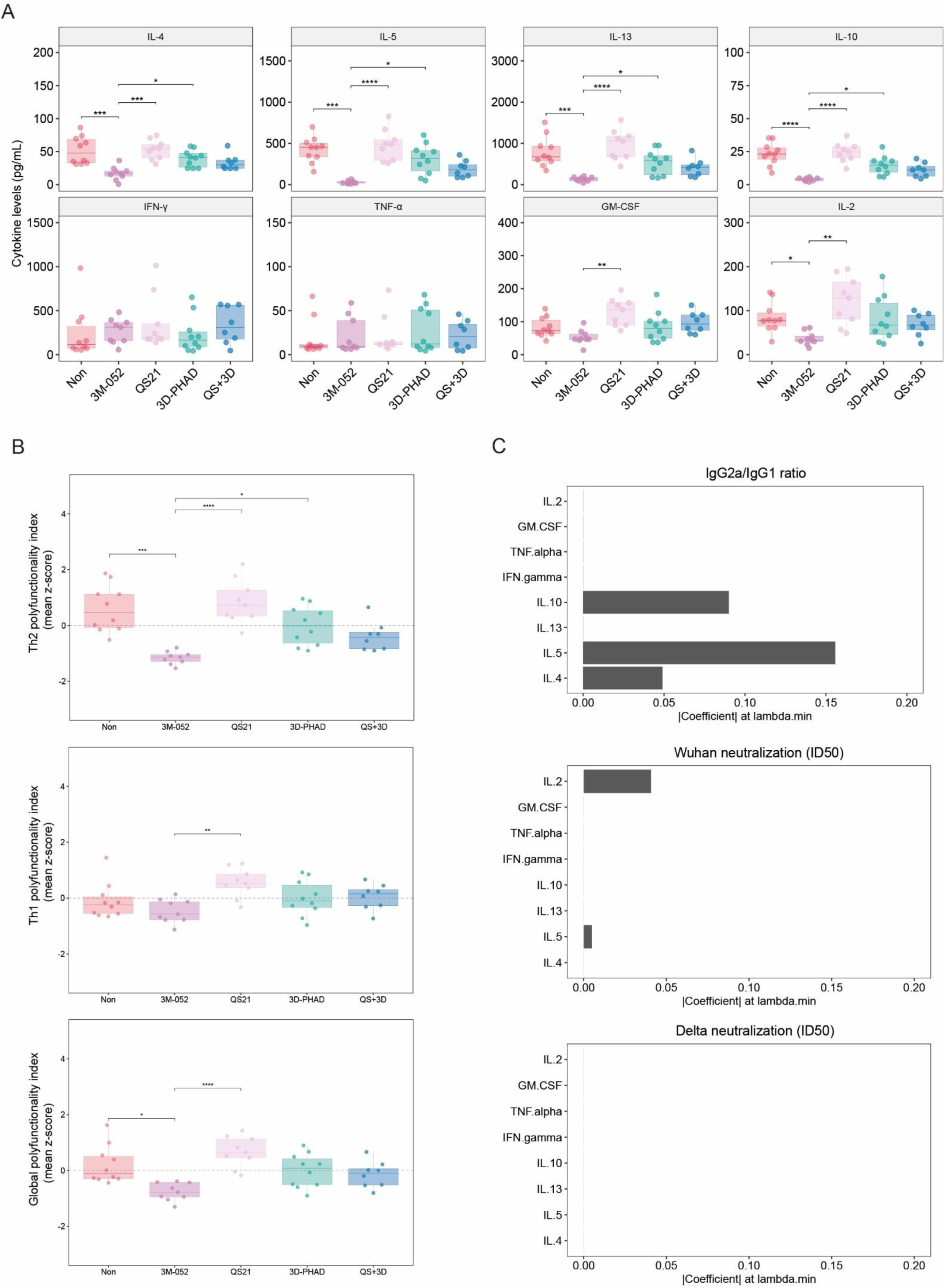
Adjuvant-dependent cytokine responses following splenocyte restimulation. Splenocytes were harvested from Wuhanvirosome immunized BALB/c mice at termination (week 13) and restimulated ex-vivo for 16 hours with a Wuhan Spike peptide pool and recombinant Spike protein. ( **A**) Cytokine secretion was quantified from culture supernatants using a Luminex multiplex assay. ( **B**) Th1-, Th2-associated and global cytokine polyfunctionality indices were calculated from summarized z-scored cytokine concentrations (**Th1**: IFN-γ, TNF, IL-2, GM-CSF; **Th2**: IL-4, IL-5, IL-13; **global**: IFN-γ, TNF, IL-2, GM-CSF, IL-4, IL-5, IL-13, IL-10). (**C**) LASSO regression identified cytokines associated with IgG2a/IgG1 ratio in plasma, Wuhan neutralization, and Delta neutralization. Bars show standardized model coefficients (non-zero features retained by LASSO). * = P < 0.05, ** = P < 0.01, *** = P < 0.001, **** = P <0.0001.

Because vaccine-induced immunity is shaped by coordinated cytokine programs rather than individual cytokines in isolation, we next evaluated overall Th1 and Th2 cytokine polarization using composite polyfunctionality indices derived from the independent cytokines secretion. Cytokines associated with Th1 responses (IFN-γ, TNF-α, GM-CSF, and IL-2) and Th2 responses (IL-4, IL-5, and IL-13) were combined into standardized functional scores, allowing integrated assessment of helper T-cell skewing across vaccine formulations. 3M-052 induced a reduced Th2 cytokine index while maintaining its Th1 cytokine profile compared to the non-adjuvanted group (Fig. 2B). In addition, 3M-052 adjuvantation induced less polyfunctional T cells compared to virosomes without adjuvants. Virosomes adjuvanted with QS21 induced a slightly higher Th1 index while maintaining its Th2 profile, thus leading to a more complex T cell response compared to non-adjuvanted mice. Th1 polyfunctionality and global polyfunctionality correlated positively with Wuhan neutralization (Fig. S2A).

To further identify which cytokines most strongly predicted IgG2a/IgG1 skewing, we performed a LASSO regression using cytokine levels as predictors and serology as outcome (Fig. 2C). The model only retained the Th2 cytokines IL-4 and IL-5, and IL-10 as important variables contributing to IgG2a/IgG1 plasma ratios. Th1 cytokines (e.g., IFN-γ, TNF-α) were not retained, suggesting that Th2 suppression, rather than absolute Th1 magnitude, is the dominant determinant of IgG2a class switching in our system. This aligns with the potent Th1-biased and Th2-suppressive profile of the 3M-052 formulation. For autologous neutralization outcome IL-2 was retained as a contributing variable, indicating that higher IL-2 levels were associated with increased Wuhan neutralization titers. In contrast, heterologous Delta neutralization was not associated with any cytokine, suggesting that none of the measured cytokines by itself are predictive of antibody maturation.

### Combined QS-21 and 3M-052 adjuvantation enhances antibody magnitude while maintaining a Th1-biased immune profile

Our findings here revealed distinct and complementary immune profiles induced by individual adjuvants, with 3M-052 promoting strong Th1-associated responses and QS-21 enhancing antibody magnitude and heterologous neutralization. It however remains unexplored whether combining these two adjuvants will have synergistic effects on the humoral and cellular immune response. To address this, we performed a second immunization experiment with the same immunization schedule using virosomes displaying the SARS-CoV-2 Delta spike protein, comparing virosomes adjuvanted with 3M-052 alone, a combination of 3M-052 and QS-21, or no adjuvant (Fig. 3A).

**Figure 3.**
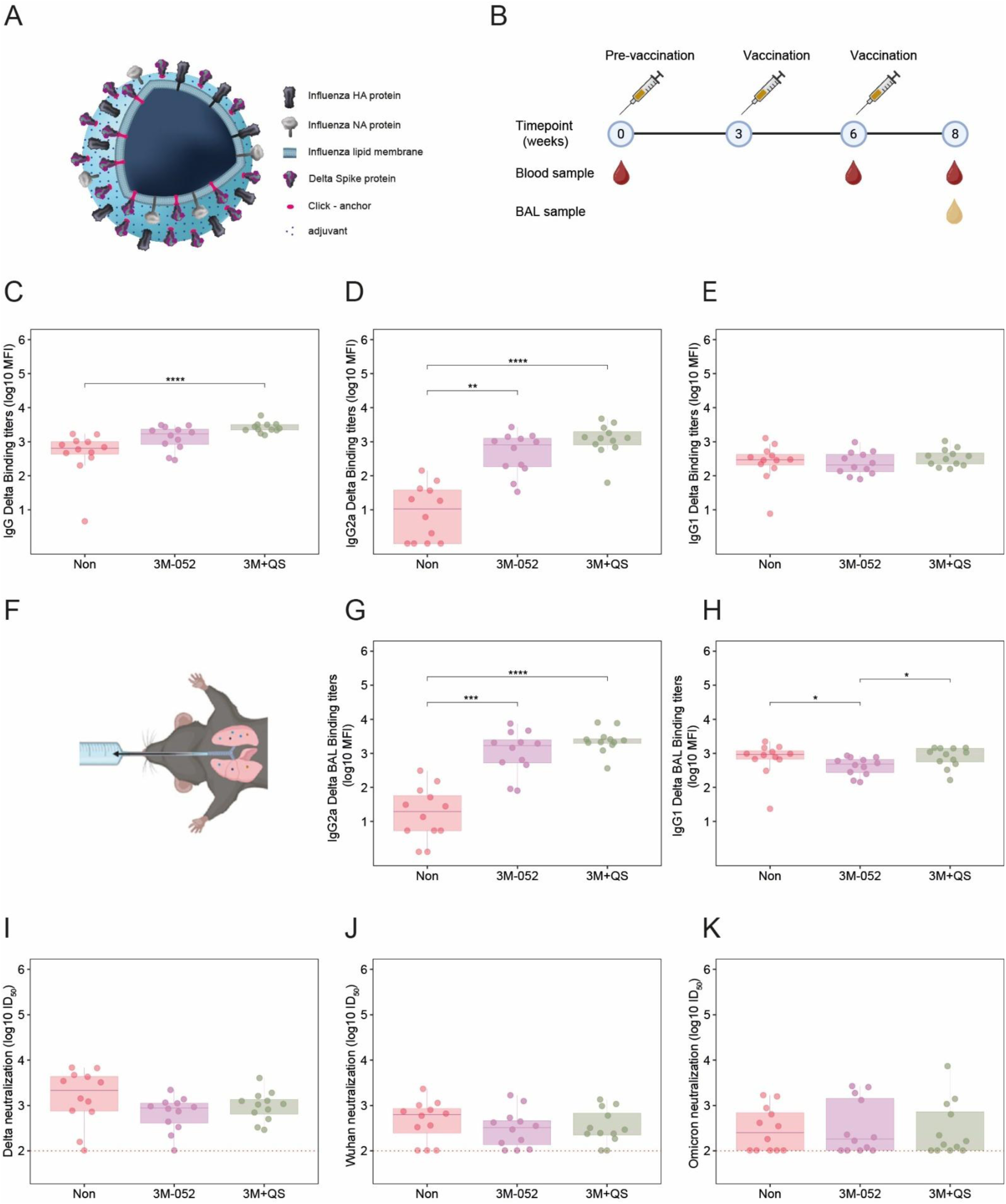
Combined adjuvant formulation has limited effect on antibody and neutralizing responses. BALB/c mice were immunized with SARS-CoV-2 Delta Spike virosomes conjugated with the following adjuvants: non-adjuvanted, 3M-052 and 3M-QS21+QS21. (**A**) An illustration of adjuvant conjugated Delta Spike virosomes. ( **B**) Serum and BAL samples were collected at indicated timepoints. ( **C-E**) Antigen-specific total IgG (**C**), IgG1 (**D**) and IgG2a (**E**) responses at week 8 were quantified for serum and for bronchoalveolar lavage (BAL) ( **F-H**) using a Luminex assay and reported as median fluorescence intensity (MFI). (**I-K**) Neutralizing activity was measured using a pseudotyped virus neutralization assay reported as ID50 against Delta (**I)**, Wuhan (**J)** and Omicron (**K)**. Data are shown for n = 12 mice per adjuvant group. Statistical comparisons were performed using the Kruskal-Wallis test with Benjamini-Hochberg (BH) correction for multiple comparisons. Dotted lines represent the lower limit of detection. * = P < 0.05, ** = P < 0.01, *** = P < 0.001, **** = P <0.0001.

Mice vaccinated with virosomes formulated with the combined 3M-052 + QS-21 adjuvant induced significantly higher plasma anti-Delta spike IgG levels compared to unadjuvanted controls ( P < 0.0001) but not compared to 3M-052 alone (Fig. 3C). Consistent with the first experiment, 3M-052 alone significantly increased anti-Delta spike IgG2a levels in plasma (P < 0.01), an effect that was further enhanced in the combined adjuvant group (P < 0.0001) (Fig. 3D). We also observed no differences in IgG1 serum levels in 3M-052 alone or combined with QS-21 compared to unadjuvanted mice (Fig. 3E). In BAL, both 3M-052-adjuvanted groups exhibited elevated anti-Delta spike IgG2a levels relative to the non-adjuvanted group (P < 0.001 for 3M-052 alone and P < 0.0001 for 3M-052 + QS-21, Fig. 3G). Interestingly, reduced IgG1 levels in BAL were observed in the 3M-052 alone group compared to non-adjuvanted mice (Fig. 3H).

Both 3M-052- and 3M-052 + QS-21-adjuvanted virosomes induced significantly higher IgG2a/IgG1 ratios in plasma compared to non-adjuvanted virosomes (P < 0.001 and P < 0.0001, respectively) (Fig. S3A). A comparable increase in IgG2a/IgG1 ratios was observed in BAL fluid (P < 0.001 and P < 0.0001, respectively) (Fig. S3B). IgG2a/IgG1 ratios were similar between the two adjuvanted groups in both compartments.

### Co-adjuvantation of QS-21 fails to enhance neutralization capacity

We next assessed whether QS-21 retained its ability to enhance neutralizing antibody responses when combined with 3M-052. Autologous neutralization against the Delta variant was unchanged in mice immunized with virosomes adjuvanted with the combined 3M-052 + QS-21 formulation compared to both non-adjuvanted virosomes and virosomes adjuvanted with 3M-052 alone (Fig. 3I). Similarly, heterologous Wuhan neutralization was unchanged in the combined adjuvant group relative to both non-adjuvanted controls and the 3M-052 group (Fig. 3J). Neutralization of the Omicron BA.1 variant was absent in most animals of all groups, with the exception of a few mice that did show neutralization (Fig. 3K).

### Addition of QS-21 slightly broadens cytokine responses without shifting Th1 dominance

QS-21 adjuvanted mice induced a polyfunctional cytokine profile. Therefore, we assessed whether combining this adjuvant with 3M-052 would recover the type 2 cytokines and increase levels of IL-2 (Fig. 4A). Similarly to the first experiment, adjuvantation with 3M-052 led to a decrease in splenic cytokine secretion of IL-4, IL-5, IL-13 and IL-10 after stimulation compared to the non-adjuvanted virosome group (p = 0.08, p < 0.001, p < 0.05, p = 0.064, respectively). QS21 + 3M-052 combined adjuvant increased the secretion of cytokines GM-CSF, IL-2 compared to 3M-052 alone, in some mice. Th2 cytokines IL-4 and IL-13 were marginally elevated compared to 3M-052 alone, in a portion of mice. The polyfunctionality index revealed a decreased Th2 immune profile in the 3M-052 and the combination adjuvanted groups compared to non-adjuvanted mice (p < 0.01 & p < 0.05, respectively) (Fig. 4B). No difference in Th1 profile and global polyfunctionality index was observed between the different groups. We performed LASSO regression analysis to further identify which cytokines most strongly predicted IgG2a/IgG1 skewing. Again the model retained IL-4, IL-5 and IL-10 as important predictors for IgG2a/IgG1 skewing (Fig. 4C). The model retained IFN-y, IL-13 and IL-4 as strong predictors for autologous Delta neutralization which is in contrast to our initial finding of IL-2 being a predictor in the first experiment. No predictors were found for heterologous neutralization of Wuhan.

**Figure 4.**
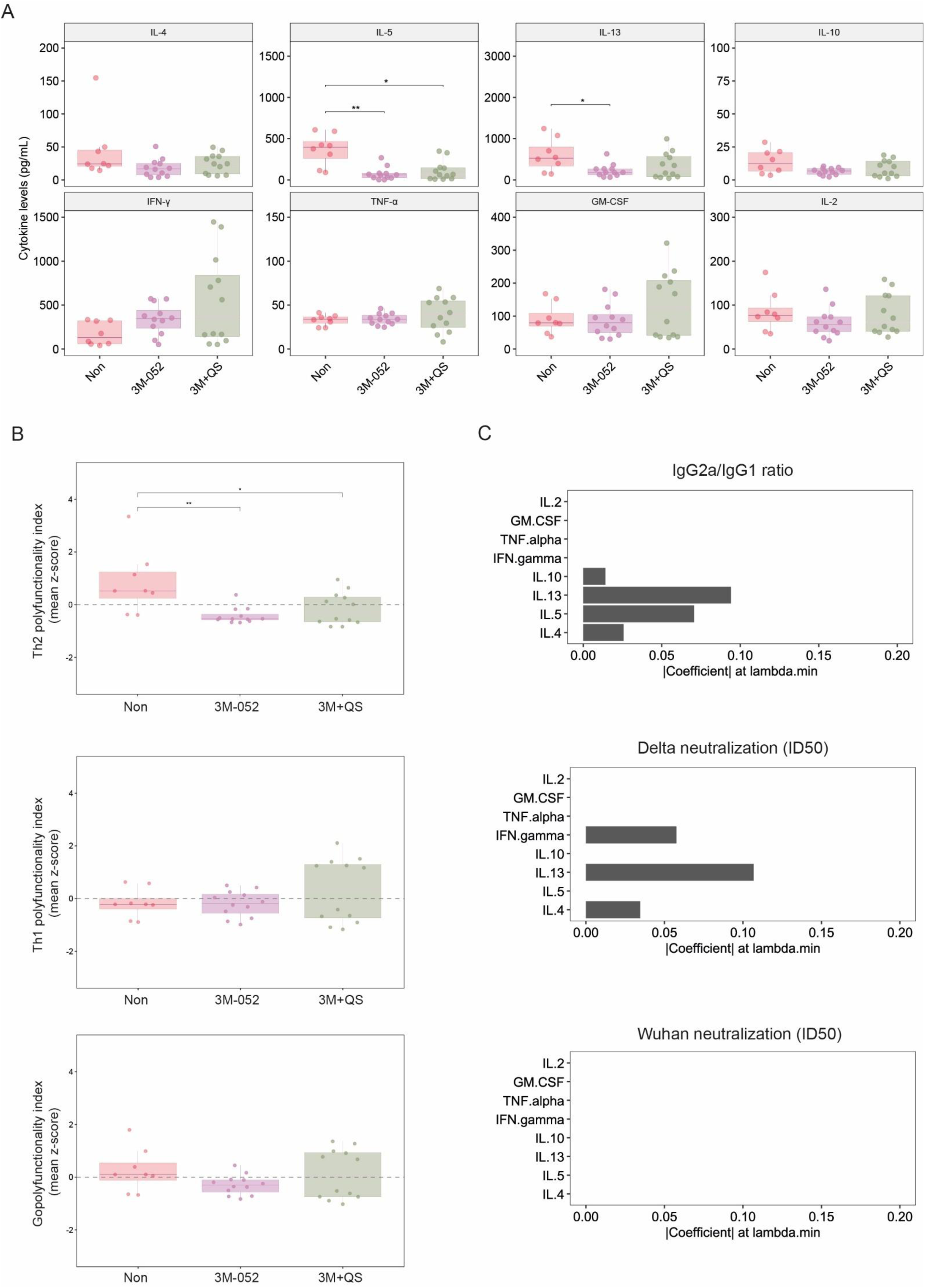
Combined adjuvant formulation fails to rescue Th2 profile. Splenocytes were harvested from Delta virosome immunized BALB/c mice at termination (week 13) and restimulated ex-vivo for 16 hours with Delta Spike peptide pool and recombinant Spike protein. ( **A**) Cytokine secretion was quantified from culture supernatants using a Luminex multiplex assay. ( **B)** Th1-,Th2-associated and global cytokine polyfunctionality indices were calculated from summarized z-scored cytokine concentrations (**Th1**: IFN-γ, TNF, IL-2, GM-CSF; **Th2**: IL-4, IL-5, IL-13; **global**: IFN-γ, TNF, IL-2, GM-CSF, IL-4, IL-5, IL-13, IL-10). (**C**) LASSO regression identified cytokines associated with IgG2a/IgG1 ratio in plasma, Delta ID50, and Wuhan ID50. Bars show standardized model coefficients (non-zero features retained by LASSO). * = P < 0.05, ** = P < 0.01.

In summary, the combination of 3M-052 and QS-21 revealed a Th1 biased antibody response similar to that of 3M-052 alone but did not significantly enhance the neutralization capacity. The addition of QS-21 failed to alter the T cell composition into a balanced Th1/Th2 profile adding to the evidence that a Th2 profile is necessary for a neutralizing response.

## Discussion

Antibody-mediated protection induced by vaccination depends on the magnitude and affinity of the response, the epitopes targeted, and antibody isotype, which together shape neutralizing breadth and potency as well as Fc-mediated effector functions. Next-generation vaccine strategies such as virosomes may aid in steering immune responses to develop these qualities through customizable adjuvant selection. Furthermore, this modular vaccine platform has previously shown promising results in licensed vaccines targeting influenza (Inflexal V, Epaxal, Nasalflu) and experimental vaccines against SARS-CoV-2, malaria, HCV, HIV and Candida (Moser et al., 2011; Van Der Velden et al., 2022). In this study, four adjuvant formulations coupled to virosomes displaying SARS-CoV-2 spike antigens were evaluated for their capacity to differentially shape humoral and cellular immune responses after immunization. Adjuvant QS-21 showed the most potent autologous and heterologous neutralization while preserving IgG isotype balance, whereas the other adjuvants did not provide additional enhancement of neutralization. The findings demonstrate that adjuvant selection leads to divergent effects on isotype polarization and neutralization potency and breadth, which is accompanied by distinct patterns of splenic cytokine responses.

In line with our findings, previous studies in non-human primates (NHPs) have demonstrated that QS-21 complexed with MPLA enhanced antibody responses through improved germinal center activity and T-follicular help support (Silva et al., 2021). Consistent with these findings, QS-21 adjuvantation in the present study selectively enhanced autologous and cross-neutralization to variant strain Delta while preserving a balanced IgG2a and IgG1 isotype distribution. TLR7/8 agonist–based adjuvants such as 3M-052 have been widely associated with Th1 polarization and IgG2a class switching, features often associated with favorable outcome for antiviral immunity (Miller et al., 2020; Smirnov et al., 2011; Vasilakos & Tomai, 2013). However, the pronounced IgG2a skewing observed here was accompanied by reduced IgG1, lower total IgG, and diminished neutralization breadth, suggesting that Th1 polarization alone does not necessarily translate into improved functional antibody outcomes. Two seminal papers on 3M-052 in NHPs showed superior antibody responses both in antibody levels and neutralization over alum-adjuvanted macaques (Kasturi et al., 2020; Pino et al., 2021). However, we observe non-superior humoral outcomes to non-adjuvanted or other adjuvant groups. One cause may be the formulation of the TLR7/8 agonist in relation to the antigen. In the present study the adjuvant is linked to the lipid bilayer of the virosome while those earlier studies used alum-absorbed 3M-052, which ultimately could alter the kinetics and activation potential of 3M-052.

Similarly, 3D-PHAD, a strong Th1 polarizing adjuvant, induced modest neutralization improvements and balanced subclass distributions, while combined adjuvants QS-21 and 3D-PHAD improved IgG2a production. This combination, however, failed to improve neutralization potency, indicating that modulation of antibody subclasses opposingly affect neutralization capacity.

Distinct patterns of splenic cytokine responses were observed, coinciding with the divergent antibody responses seen across different adjuvant formulations. QS-21 elicited a more polyfunctional and stronger in magnitude T cell response characterized by the induction of Th1 cytokines such as IFN-γ, TNF-α, GM-CSF and IL-2 while preserving Th2 cytokines such as IL-4, IL-5 and IL-13. In contrast, 3M-052 elicited a diminished Th2 cytokine response, indicating a unidirectional polarization towards a Th1 response. These differences suggest that a balanced Th2 and Th1 cytokine profile may better support (maturity of) the humoral response. Such balanced T cell help may enable antibody quality and breadth through coordinated support of B cell differentiation and selection processes, whereas strongly polarized environments may constrain functional outcomes despite pronounced inflammatory signatures.

These findings further raised the question of whether combining adjuvants with distinct immunological profiles could integrate complementary immune features within a single formulation. Combining 3M-052 (TRL7/8 agonist) with MPLA (TRL4 agonist) exhibited greater humoral responses in NHPs (Silva et al., 2021). Here, combining 3M-052 and QS-21 still resulted in Th1 skewed responses and an antibody profile shifted toward IgG2a while diminishing IgG1 responses, and failed to enhance either autologous or heterologous cross-neutralization relative to non-adjuvanted immunization. This outcome suggests that the strong Th1 context imposed by TLR7/8 agonism, such as 3M-052, may dominate immune programming, limiting the functional benefits conferred by the more balanced immune modulation associated with QS-21. Together, these observations indicate that simultaneous engagement of multiple immune pathways does not necessarily yield additive or synergistic improvements in antibody functionality.

This study was not designed to directly assess germinal center dynamics or B cell receptor maturation which limits our mechanistic understanding of how these adjuvants influence isotype switching and neutralization breadth. In addition, regression-based models used here identify correlates rather than causal drivers of antibody outcomes. Moreover, one key cytokine associated with humoral help, IL-21, was not studied here, limiting our understanding of the contributions of Tfh/Tfh-like cells. Further experimental validation will be required to elucidate the mechanistic relationship between these cytokines and humoral outcomes. Finally, these findings were generated in BALB/c mice which are inherently Th2 biased and may overestimate humoral responses. While this model is well suited for evaluating antibody induction and subclass distribution, validation in additional genetic backgrounds, such as C57BL/6 mice, will be important to assess the generalizability of the observed adjuvant effects.

Collectively, the findings here support a model in which antibody magnitude, function and neutralization capacity emerge from a balanced immune programming rather than a Th1 skewed polarization. Adjuvants that preserve balanced T helper cell activity may better support antibody diversification and selection processes that favor neutralization potency and breadth. In this context, QS-21–induced immune responses appear to integrate multiple T helper cell signals that support functional antibody maturation, whereas strongly polarized environments driven by 3M-052 favor subclass skewing towards IgG2a without conferring comparable gains in neutralization capacity. These findings may help guide rational vaccine design by tailoring adjuvant strategies that promote a balanced humoral and cellular programming for viral pathogens in the context of virosomes.

## Methods

### Protein design

Pre-fusion spike protein ectodomain DNA constructs corresponding to the Wuhan (Wuhan-Hu-1; GenBank MN908947.3) and Delta variants were designed and ordered as gene fragments (Integrated DNA Technologies). The constructs were inserted into a pPPI4 expression vector containing a hexahistidine (His) tag using Gibson Assembly (ThermoFisher). All spike constructs were produced in HEK293F cells (ThermoFisher) and purified using NiNTA chromatography and size exclusion chromatography as previously described (Brouwer et al., 2020).

### Virosome generation

Virosomes were generated from detergent-solubilized influenza virus membranes of which lipid head-groups were modified for click chemistry (DBCO-PE) as previously described (Amacker et al., 2020). Briefly, inactivated influenza A/Brisbane/59/2007 was solubilized with OEG, and the viral nucleocapsid was removed by centrifugation. Dioleoyl-phosphatidylcholine, cholesterol, and DBCO-PE were then added to the supernatant, after which OEG was removed using BioBeads SM2 and the virosomes were sterilized by filtration. ATPD was synthesized and purified as described previously. The S protein was dialyzed against 50 mM HEPES buffer (pH 8.5), reacted with ATPD at a 200:1 ratio for 1 h at room temperature, and dialyzed overnight against buffer containing NaCl, HEPES, and EDTA. The resulting azide - modified S protein was filter-sterilized and incubated with virosomes for at least 24 h at 25 °C to allow covalent coupling through azide-DBCO click chemistry. S protein concentration was estimated by SDS-PAGE. The adjuvantswere incorporated into the virosomal membrane by post-insertion, and coupling of S to virosomes was confirmed by ELISA.

### Mouse immunization

Female BALB/cAnNCrl mice were vaccinated with virosomes as previously described (Van Der Velden et al., 2022). Briefly, mice were first given a subcutaneous pre-vaccination with inactivated influenza A/Brisbane/59/2007 in the neck skin fold at week 0 to model pre-existing influenza immunity. At weeks 3 and 6, they were immunized subcutaneously in the same site with virosomes containing adjuvanted Wuhan or Delta SARS-CoV-2 spike protein, with 16 mice included in each group. Blood samples were collected at weeks 0, 6, 8, and 13. At week 8, ten mice per group were euthanized, while the remaining six were euthanized at week 13 to assess response durability. Animals were housed under BSL-2 conditions at the Animal Research Institute Amsterdam. All experiments were conducted in compliance with the Dutch Experiments on Animals Act, approved by the Animal Ethics Committee of Amsterdam UMC (permit #202011565), and performed in accordance with ARRIVE guidelines.

### Serology assays

An inhouse luminex assay was used as described previously (Grobben et al., 2025). Briefly, spike proteins were covalently coupled to Luminex MagPlex beads using a two-step carbodiimide reaction at a ratio of 75 µg protein per 12.5 million beads. Based on prior optimization, plasma samples were diluted 1:50,000 and BAL samples 1:10. Beads were incubated overnight with diluted samples, and bound antibodies were detected using goat anti-mouse IgG-PE. Measurements were acquired on a Magpix instrument (Luminex). Mean fluorescence intensity (MFI) values were calculated as the median signal of approximately 50 beads per well after subtraction of background valuesfrom buffer-and bead-only control wells.

### Pseudovirus neutralization assay

Pseudovirus neutralization assays were performed as previously described (Caniels et al., 2021). In brief, HEK293T cells expressing ACE2 were seeded in poly-L-lysine-coated 96-well plates. The next day, heat-inactivated serum samples were serially diluted in triplicate, mixed 1:1 with SARS-CoV-2 pseudovirus, incubated for 1 h at 37 °C, and subsequently added to the cells at a 1:1 ratio. Each pseudovirus variant was used at a titer of 1000 TCID50. After 48 h, cells were lysed and transferred to half-area 96-well white microplates (Greiner Bio-One), and luciferase activity was measured using the Nano-Glo Luciferase Assay System (Promega) on a Glomax instrument (Turner Biosystems). Relative luminescence units were normalized to cells infected with pseudovirus in the absence of serum. Neutralization titers (ID50) were defined as the serum dilution that resulted in 50% inhibition of infectivity.

### Multiplex cytokine analysis of stimulated splenocytes

Mouse splenocytes were isolated as single-cell suspensions, subjected to red blood cell lysis, and resuspended in complete RPMI 1640 medium. Cells were plated in triplicate at 2.0 × 10^5 cells per well in 96-well round-bottom plates rested for a minimum of 2h and stimulated afterwards with Wuhan or Delta Spike antigen at 9.3 nM for 72 hours at 37 °C and 5% CO2, whereas medium alone and Phorbol 12-myristate 13-acetate (PMA) stimulation were included as negative and positive controls, respectively. After stimulation, supernatants were collected and analyzed using a mouse multiplex bead-based cytokine assay (R&D systems cat# LXSAMSM) on a Luminex instrument according to the manufacturer’s instructions. Cytokine concentrations were determined from standard curves and corrected for background by subtraction of values obtained in unstimulated control wells. Cytokine polyfunctionality indices were calculated by first standardizing individual cytokine measurements across samples using z-score normalization. Cytokines were then grouped according to functional profiles (Th1: IFN-γ, TNF-α, GM-CSF, and IL-2; Th2: IL-4, IL-5, and IL-13). For each sample, the polyfunctionality index was defined as the mean z-score of all cytokines within the corresponding functional group. A global polyfunctionality index was additionally calculated using the mean z-score across all cytokines analyzed

### Data analysis

Data were analyzed in R (v4.5.1) using RStudio (2025.05.1 Build 513). Statistical comparisons between groups were performed using the Kruskal–Wallis rank-sum test followed by pairwise Dunn’s tests with Benjamini–Hochberg (BH) correction for multiple comparisons. Adjusted *P* values < 0.05 were considered statistically significant. Data are presented as log10-transformed median ± interquartile range (IQR), unless otherwise indicated.

---

**Figure S1.**
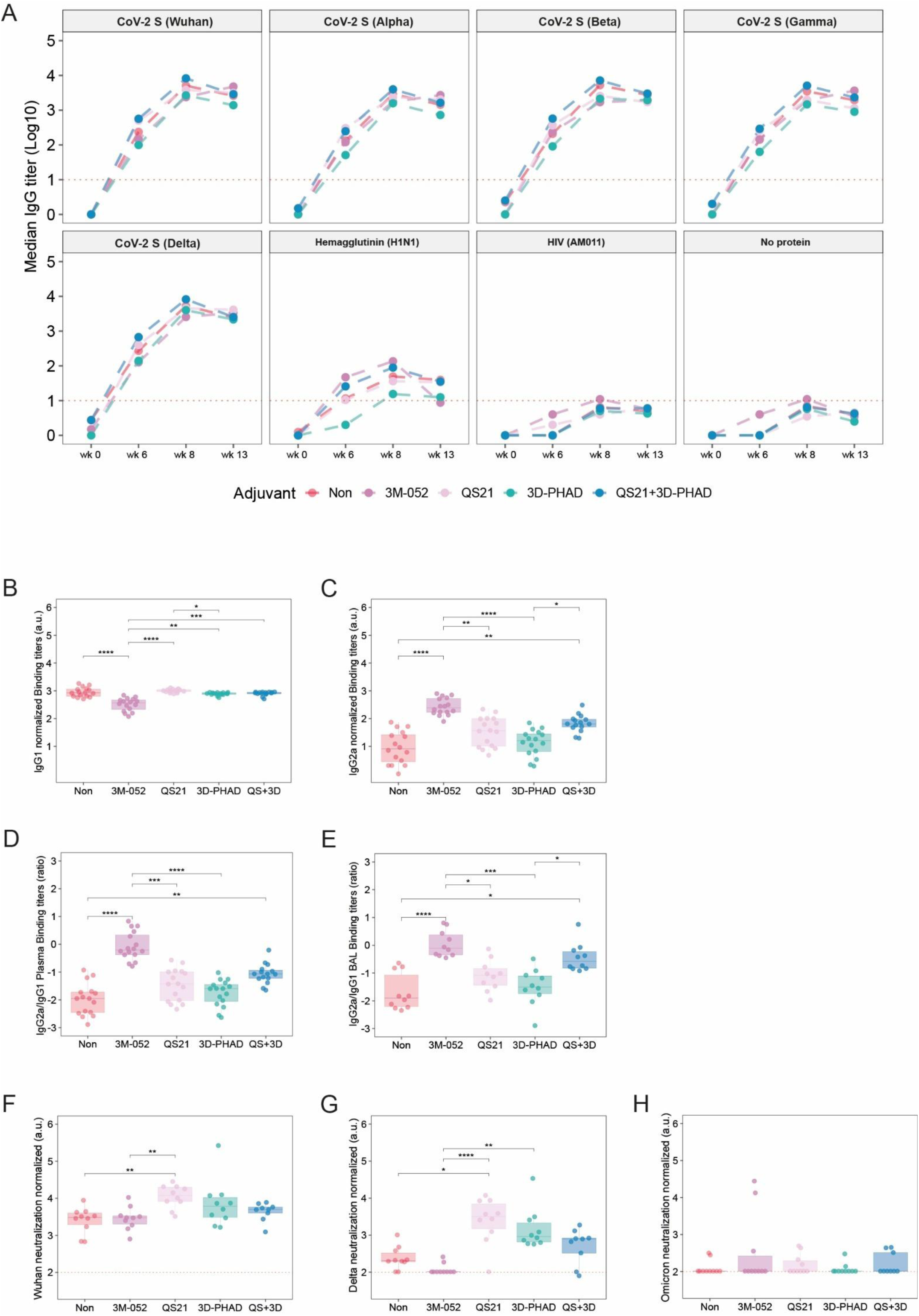
Longitudinal antibody kinetics and normalization. Serum antibody responses were measured at week 0, 6, 8 and 13 using Luminex. ( **A**) Antigen-specific binding was assessed against SARS-CoV-2 antigens corresponding to Wuhan, UK, SA, BR, IN, Influenza HA (H1N1/2009) and HIV Env (AMC011) included as control antigens and a no-protein condition included as a negative control. (**B, C**) IgG subclassing skewing independent of total IgG were expressed as a proportion of total antigen-specific IgG, IgG1/IgG (**B**) and IgG2a/IgG (**C**). (**D, E**) Th1/Th2 skewing was further calculated as IgG2a/IgG1 ratio in plasma ( **D**) and BAL (**E**). (**F-H**) Functional neutralization was assessed by normalizing ID50 over antigen-specific IgG binding titers of the corresponding adjuvant groups against Wuhan (**F**), Delta (**G**) and Omicron (**H**). Data are shown for n = 16 mice per adjuvant group. Statistical comparisons were performed using the Kruskal-Wallis test with Benjamini-Hochberg (BH) correction for multiple comparisons. Dotted lines represent the lower limit of detection. A.u. = arbitrary units, * = P < 0.05, ** = P < 0.01, *** = P < 0.001, **** = P <0.0001.

**Figure S2.**
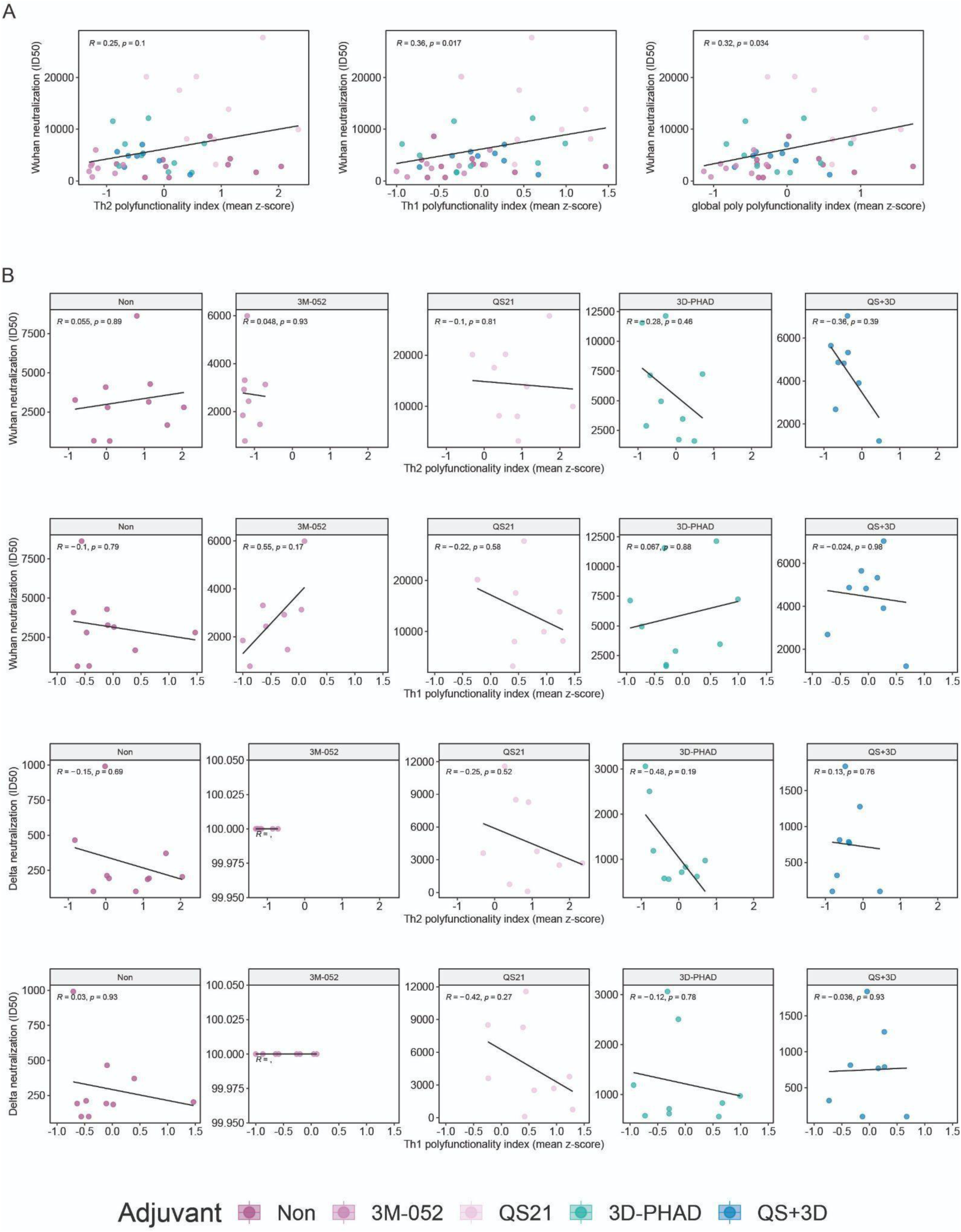
Correlations between cytokine polyfunctionality and Wuhan neutralization. (**A, B**) Associations between Th1, Th2, and global cytokine polyfunctionality indices and Wuhan pseudovirus neutralization titers (ID50) were assessed across immunized mice ( **A**) or stratified by adjuvant (**B**). Correlations were calculated using Spearman’s rank correlation. Each point represents an individual mouse.

**Figure S3.**
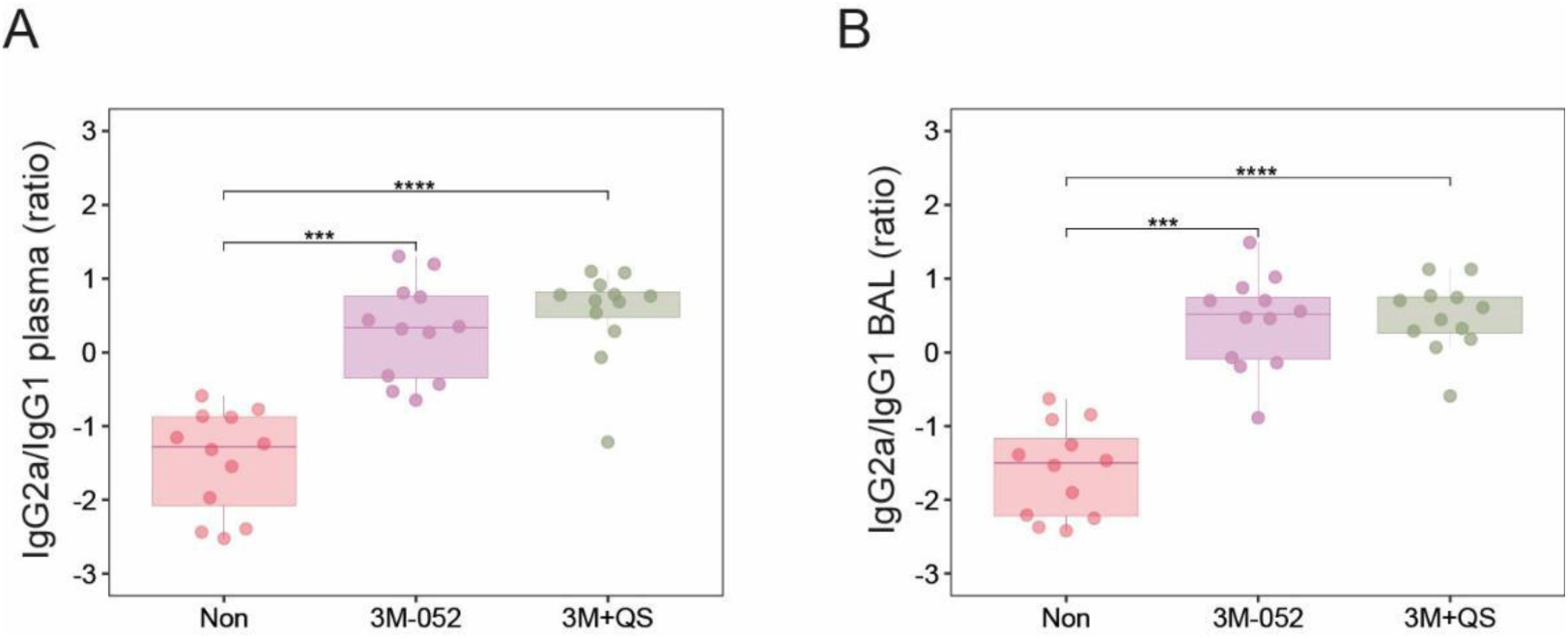
Th1 skewed antibody responses. Th1/Th2 skewing was calculated as IgG2a/IgG1 ratio in (**A**) plasma and (**B)** bronchoalveolar lavage (BAL). Data are shown for n = 12 mice per adjuvant group. Statistical comparisons were performed using the Kruskal-Wallis test with Benjamini-Hochberg (BH) correction for multiple comparisons. *** = P < 0.001, **** = P <0.0001.

